# A fragment-based protein interface design algorithm for symmetric assemblies

**DOI:** 10.1101/2021.01.13.426605

**Authors:** Joshua Laniado, Kyle Meador, Todd O. Yeates

## Abstract

Theoretical and experimental advances in protein engineering have led to the creation of precisely defined, novel protein assemblies of great size and complexity, with diverse applications. One powerful approach involves designing a new attachment or binding interface between two simpler symmetric oligomeric protein components. The required methods of design, which present both similarities and key differences compared to problems in protein docking, remain challenging, and are not yet routine. With the aim of more fully enabling this emerging area of protein material engineering, we developed a computer program, Nanohedra, to introduce two key advances. First, we encoded in the program the construction rules (*i*.*e*. the search space parameters) that underlie all possible symmetric material constructions. Second, we developed algorithms for rapidly identifying favorable docking/interface arrangements based on tabulations of empirical patterns of known protein fragment-pair associations. As a result, the candidate poses that Nanohedra generates for subsequent amino acid interface design appear highly native-like (at the protein backbone level), while simultaneously conforming to the exacting requirements for symmetry-based assembly. A retrospective computational analysis of successful vs failed experimental studies supports the expectation that this should improve the success rate for this challenging area of protein engineering.

## INTRODUCTION

A range of emerging bionanotechnology applications rely on designing protein molecules to bind and associate with each other in a geometrically specific fashion. Among such applications, those aimed at creating novel, self-assembling symmetric architectures, such as protein cages and extended protein arrays, place especially strict demands on achieving atomically precise associations [1]. When such precision can be achieved by design, diverse protein-based materials with tailored spatial and biochemical properties can be produced. As examples, cubic and icosahedral protein cages [2–7], as well as extended protein arrays [8–11], are finding wide ranging uses as biotherapeutics (e.g. for vaccines) [12–14], as scaffolds for enzyme organization or atomic imaging [15–18], and as nanoscale containers for molecular encapsulation and delivery [19,20].

Owing to their complexity, as well as our incomplete understanding of their behavior, protein molecules present challenging subjects for design. In protein engineering studies, these challenges often manifest through unpredictable outcomes from mutagenesis, frequently leading to proteins that are prone to misfolding and aggregation. Improved computational methods are addressing those challenges, making it increasingly feasible to mutate the surface of two suitably chosen proteins to create a binding interface between them [21–23]. Similar goals are being reached using *de novo* polypeptides as components [24–27]. Yet, despite exciting progress, relatively low success rates are still common in application areas where precision and predictability are essential, generally requiring many design trials to achieve a smaller number of correctly assembling protein designs.

A general view on the current challenges in designing novel protein-protein interfaces is that computational methods do not necessarily generate (prospective) interfaces that mimic native protein-protein interfaces [28]. The difficulty of the task is heightened in design problems where additional spatial constraints must be met, beyond those required simply for binding. For the design of symmetric cages and regular arrays, for instance, the novel interface must bring the two component proteins together under exacting rules of symmetry; e.g., if each component is part of a naturally symmetric oligomer, then the interface must cause the symmetry axes of the separate components to intersect at a precisely prescribed angle. Such complex constraints confound the problem of designing optimal, native-like interfaces.

In addressing the problem of interface design in the context of symmetric assembly, the strategy introduced by King [3] prioritized the symmetric constraint part of the problem. There, oligomeric building blocks were docked by systematically sampling the rigid body degrees of freedom allowed by the point symmetry of the target assembly. As a result of the high dimensionality search space and the large number of different component oligomers considered for docking, a rapid first-pass scoring was used to identify configurations that were potentially suitable for design: the number of Cβ contacts between the docked oligomeric building blocks. Naturally, only a minute fraction of candidate poses chosen under such coarse criteria present interfaces that are similar in atomic detail to those from natural protein-protein complexes. Subsequent amino acid sequence design and additional filtering steps were required to identify interfaces that might exhibit native-like properties. Newer protocols have shown the value of considering known residue pair interactions during docking [22] and prioritizing interfacial hydrogen bonding during sequence design [6,25,29].

The expansive database of known protein structures provides valuable empirical frameworks for evaluating proteins in terms of secondary structure motifs [30–34]. Recent exercises in protein design have begun to prioritize the consideration of secondary structure motifs and the atomic details of how they tend to associate in native proteins [35,36]. For instance, threading helical fragments together produces novel fold topologies that retain features of observed tertiary motifs [37,38]. Further, sequence design using statistical models of tertiary structure segments has competed with or outperformed physicochemical energy functions in routine design tasks [39]. The growing focus on secondary structure associations motivates an attempt to bring those principles to bear on the class of design problems related to symmetry-based assemblies.

Here we describe algorithms and software that expand motif-based design methodologies to symmetric docking applications – e.g. cubic cages and extended protein arrays. Our new program is parameterized to exploit recent theoretical work articulating the geometric rules for designing wide ranging nanoscale materials built from combinations of oligomeric protein components – i.e. symmetry combination materials (SCMs) [40]. Strategic choices are discussed for program optimization based on fragment-based lookup tables and separation of rotational vs translational subspace searches. Prospective novel designs are discussed, along with a retrospective analysis of successfully designed protein cages.

## RESULTS

### Docking under symmetry constraints

The goal of the program developed here was to enable fragment-based docking for the design of self-assembling materials based on the principles of combined symmetries. The essential idea for building highly symmetric materials from simpler protein oligomers was described by Padilla et al. [2], with diverse variations demonstrated in recent years [4,5,7,41]. A complete articulation of all possible SCMs was recently completed [40]; 124 different kinds of architectures can be created by introducing a new interface between two oligomeric components. In addition to various cage types based on the Platonic solids, 35 kinds of 2-D arrays and 76 kinds of 3-D arrays were identified as targets possible for design. Each of the distinct SCMs presents a different set of rigid body constraints, and complementary rigid body degrees of freedom, for sampling allowable arrangements of the two oligomers to be docked. For the present work, we have integrated the design rules for all possible SCMs within a new program, Nanohedra.

We developed a general docking framework, applicable to all SCMs, that performs a search over multiple rigid body degrees of freedom relating two oligomeric building blocks (Fig. 1). The number of degrees of freedom depends on the symmetric system being constructed, ranging from a minimum of 1 to a maximum of 5 [40]. Exploiting advantages of pre-calculation methods, we were able to factor the search problem for all scenarios into a search over rotational degrees of freedom (for cases where they exist), followed by direct calculation of optimal translational values by linear algebra methods, thereby avoiding the need to explicitly search translational degrees of freedom for each rotation. Identifying favorable docking arrangements within the allowable rigid body search space is made possible by precomputing common protein-protein fragment configurations from known structural data.

**Fig. 1.**
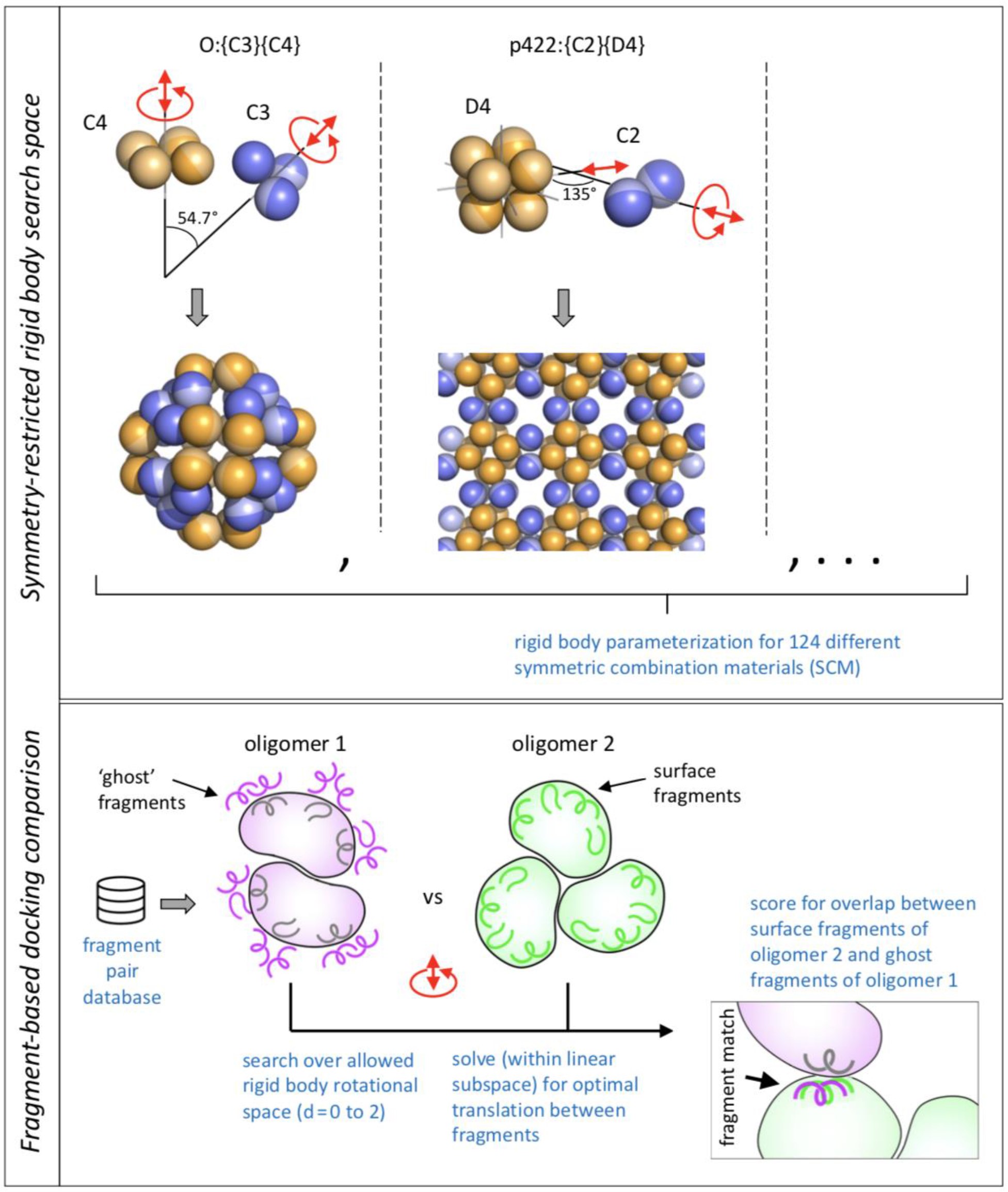
Scheme illustrating two major aspects of the Nanohedra program for designing symmetry combination materials (SCMs) from two oligomeric protein components. The top panel shows examples of two SCM types (of 124 types possible), focusing on the geometric rules that must be satisfied when bringing the two different oligomeric components into specific contact. In each case, the red arrows indicate the rigid body degrees of freedom available, which must be explored computationally in a search for favorable docking configurations that would be amenable to amino acid sequence design at the emergent protein-protein interface. Nanohedra encodes the specific rigid body parameterization required for constructing all 124 SCM types [40]. The bottom panel highlights the use of protein fragment pair libraries as the essential feature for selecting favorable design poses for subsequent interface design. This allows Nanohedra to generate native-like interfacial backbone arrangements for design. Program operation is made computationally tractable through various pre-calculation schemes. One of these involves the decoration of the first oligomer with ‘ghost fragments’ (based on a library of favorable fragment pair configurations), after which the search for suitable docking poses is reduced to a problem of identifying allowable oligomeric arrangements wherein surface fragments belonging to oligomer 2 overlap closely with ghost fragments covering oligomer 1.

### Fragment-based elements

Focusing on short segments of protein structure makes it possible to reduce computational complexity with lookup or ‘hash’ tables. To this end, we chose to categorize local protein structure using 5-residue fragments. Heuristically, a five-residue segment is long enough to capture secondary structures types, including α-helical and β-strand conformations, as well as loop structures, while being short enough to model the allowable space of conformations with acceptable precision and coverage using a tractable number of representatives. Using a curated set of known protein-protein interfaces (see Methods), we computed the most highly represented 5-residue fragment types found at interfaces using nearest neighbor clustering (on Cα RMSD) for a randomly sampled subset of fragments. We experimented with different similarity cutoff criteria, and settled on a 0.75 Å cluster inclusion limit, which maximized fragment coverage, while ensuring stringent constraints on backbone geometry.

As expected, different fragment clusters were populated to different degrees; those representing canonical α-helical and β-strand conformations were much more densely populated than those representing different loop conformations, or cases where regular secondary structure transitioned into loops. The top five clusters were sufficient to represent 61.4% of the candidate fragments; with the highest observed cluster corresponding to an α-helical conformation, followed by a β-strand conformation then three coiled conformations (Fig 2). Rather than considering a larger number of clustered fragment conformations, we retained five cluster types in order to maximize statistical power in subsequent steps.

**Fig. 2.**
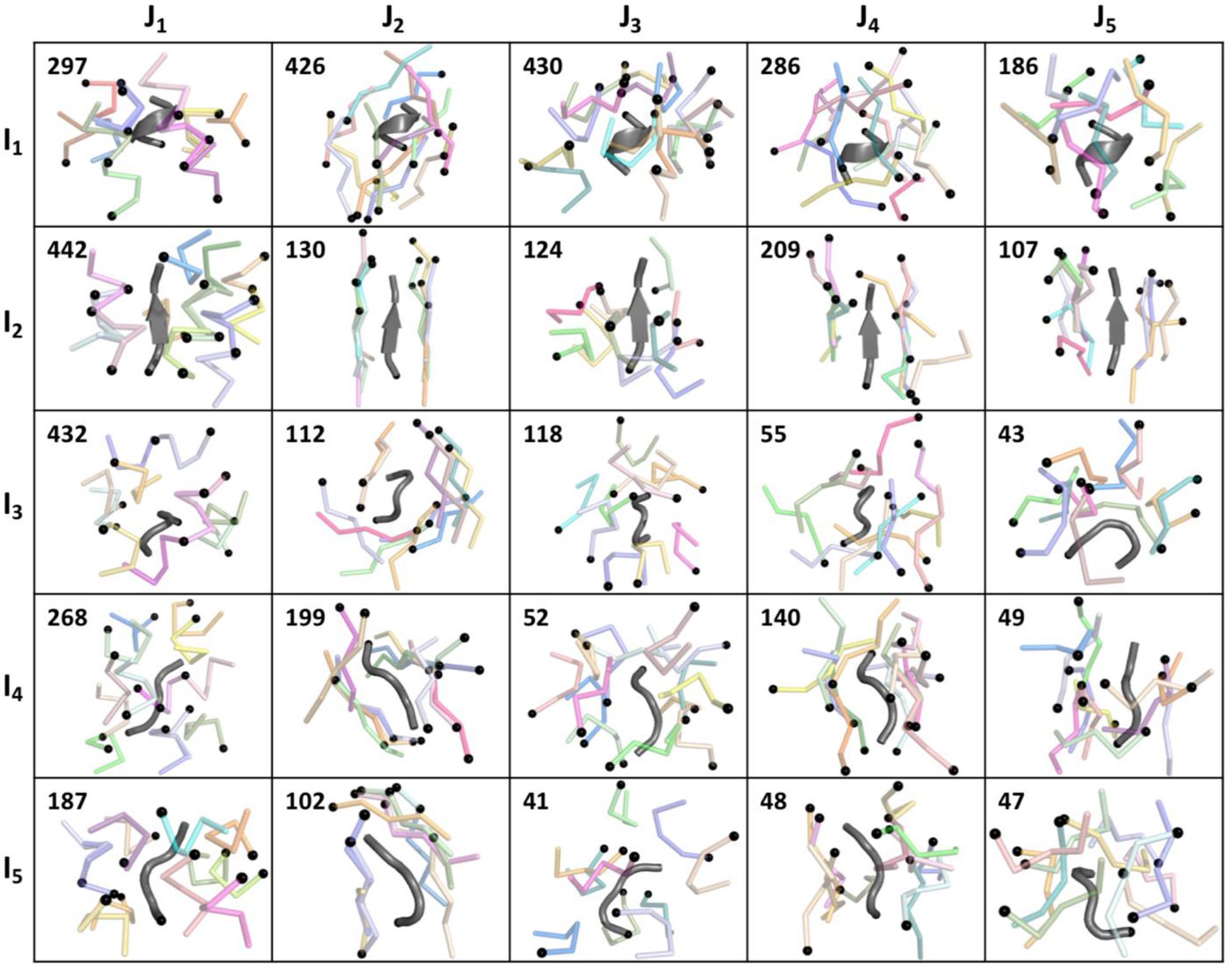
Interface fragment database. For each of the 25 *i,j* fragment pair possibilities, a single representative *i* fragment is shown in grey and a subset of cluster representatives of the top 20 most populated clusters are shown in color for the spatially clustered *j* fragments. N-termini are marked with black spheres. The total number of unique *i,j* clusters is indicated in the top left corner of each frame.

Following individual fragment clustering, a paired fragment-fragment clustering procedure was applied to inter-protein contacts from the same protein-protein interface set. This problem was simplified by a form of coordinate reduction. A set of three ‘guide coordinates’, built on the C-alpha atom of the central residue of the 5-residue fragment (Fig. S1), was associated with the representative fragment from each individual fragment cluster (see Methods); note that three x,y,z coordinates (nine variables) are sufficient to specify six dimensional rigid body orientation and position in three-dimensional space. This provided a generalized scheme for clustering the relative spatial arrangement between fragment-fragment pairs. Briefly, for every instance where a 5-residue fragment (type *i*) from one protein was found in spatial contact with a 5-residue fragment (type *j*) from another protein, each fragment in the *i,j* pair is assigned to one of the five individual fragment types. This allows placement of the representatives’ guide coordinates onto the coordinate frame of the observed fragment pair. Next, the guide coordinate pair was transformed to put the *i* guide coordinate set in a canonical setting (i.e. at the origin with internal axes along principle directions). The resulting *j* guide coordinate set is then stored, providing a full representation of the relative spatial arrangement of the *i,j* fragment pair instance. The *j* guide coordinate sets were then used as the basis for a final nearest neighbors clustering step where the resulting cluster index, *k*, represents the different spatial modes that tend to be populated by specific *i,j* fragment pairs (Fig. 2). Clustering at this pairwise stage was based on a relatively strict similarity criterion (1 Å guide coordinate RMSD) to establish separate conformational and amino acid preferences for relatively finely discriminated fragment-fragment arrangements (Fig. S1). The resulting data structure is a triplet of (*i,j,k*) indices, each carrying a 9-dimensional coordinate point that captures a frequently observed spatial relationship *k*, between a specific *i,j* fragment pair type. In addition, owing to the cartesian nature of the embedding, a 9×9 covariance matrix (approximately rank 6) provides a quadratic description of the spatial variation for the given *i,j,k* fragment pair cluster; that information can be used to analyze permissible deviations. Observed central residue amino acid frequencies are also stored for each *i,j,k* cluster and can be used to guide subsequent sequence design steps. Ultimately, a total of 97,935 *i,j* fragment pairs from observed structures were grouped into 4,530 *i,j,k* clusters specifying geometrically defined 3-D fragment associations.

### Precoating by ‘ghost fragments’

Having established common fragment pairing conformations in advance enables a pre-calculation protocol with important computational time savings. As a set-up to docking trials between two oligomers, one of the component oligomers (the first oligomer) is decorated with a large set of prospective ‘ghost fragments’ (Fig. S2 and Methods). These ghost fragments represent preferred interaction potentials based on the orientation of the fragments on the surface of the first oligomer. The precalculated database of representative *i,j,k* fragment pairs described previously serves as the source for constructing this set of ghost fragments, which is intended to be inclusive for the backbone configurations of the second oligomer that might comprise frequently observed interactions with the first. Depending on the size of the first protein, the ghost fragments may number in the thousands. Once the ghost fragment set is calculated, the subsequent fragment-based docking scheme is reduced to a problem of identifying orientations that might bring surface fragments of the second oligomer into near coincidence with the ghost fragments decorating the first.

### Identifying favorable rotations and translations

The outer loop of the docking calculations applies candidate rotation values to the two oligomers, if those degrees of freedom exist. The symmetric construction schemes for SCMs provide a maximum of one degree of rotational freedom for each component oligomer (e.g. about the unique symmetry axis of a cyclic oligomer). For each choice of rotational values for the two oligomers, a calculation is performed to test which of the possible pairs of fragments (chosen from the surface fragments of 2 and the ghost fragments of 1) are in nearly equivalent orientations, as would be required for near-overlap under any choice of translation. This step involves a large number of possible fragment pairs to be considered, as well as somewhat complex numerical calculations for orientation comparisons. We found it critical to shorten this calculation with further pre-calculation methods and hash tables. We assign each fragment (based on its guide coordinates) to a set of three Euler angles describing its orientation, with the Euler angles discretized into 10° bins. With a triplet of orientation indices assigned to each fragment, we are able to look up in a precalculated 6-dimensional Boolean (true/false) table whether or not the sets of triplets assigned to the two fragments are within a prescribed angular discrepancy (with an accuracy of roughly 10°).

The steps described above rapidly identify pairs of fragments (a surface fragment of oligomer 2 and a ghost fragment surrounding oligomer 1) that could be nearly coincident under the chosen orientation values and *some* translational values between the oligomers. It is critical however that the translational relationships conform to those that are prescribed by the particular symmetry rules of the SCM being constructed. Some SCM types have three translational degrees of freedom while some have as few as one. Importantly, our program encodes those translational restrictions for all SCM types, based on tables provided in Laniado and Yeates [40]. For every pair of candidate fragments that have compatible orientations, our program calculates the optimal translation for overlap, *within the allowable space of rigid body translations for the given SCM type*. This is performed using a linear least squares calculation, with the error value based on rms deviation between the two sets of guide coordinates as a function of translational degrees of freedom. We then store the translational parameters for cases where the RMSD for the optimal overlap is within a prescribed cutoff (e.g. 1 Å).

Ultimately, a suitable docking arrangement between the oligomers is one where multiple candidate fragment pairs could be brought into near coincidence for the same (or highly similar) choices of the rotational and translational parameters. For fastest performance we found it efficacious to perform the docking analysis in a rapid first pass over a reduced set of candidate fragment pairs (e.g. requiring at least one helix-helix association), followed by a second pass wherein the translational values established in the first pass serve to restrict consideration of additional fragment pairs in the second pass, with an attendant reduction in CPU time.

We found the procedures described above critical for reducing the CPU times to levels that were compatible with docking large sets of candidate oligomer pairs. Other approaches could also be considered, though we emphasize that procedures that might appear beneficial for certain kinds of symmetric construction choices are sometimes problematic for other types of constructions, e.g. depending on the types and numbers of the rigid body degrees of freedom. The system we developed applies universally to all 124 SCM types.

### Heuristic scoring

For each satisfactory docking configuration, a Nanohedra score is calculated based on the collection of favorable fragment pairs identified, with the goal of evaluating how well the docked interface is supported by the underlying fragment observations. To compute the Nanohedra score, for each instance where a favorable surface fragment *vs* ghost fragment pair has been identified, a similarity score (z) is first calculated by dividing the RMSD obtained between the surface and ghost fragments by the mean RMSD for member fragments comprising the ghost fragment’s i,j,k cluster (precalculated during fragment database creation), with a low value of z indicating a close similarity. If z is less than a prescribed threshold value (e.g. 2), the inverse of 1 plus z squared is taken to give a match score, ranging from 0 to 1 with 1 indicating a perfect match. This match score for each fragment is propagated to each of the five residues comprising the fragments on oligomers 1 and 2. In this way, each residue in the protein-protein interface might inherit multiple component scores, since each residue might belong to overlapping fragments participating in favorable fragment-fragment pairs. For each such interfacial residue, its assigned match score(s) are first ranked in descending order and are then weighted by 1/2^rank– 1^ (rank > 0) for a final summation. This weighting scheme bounds the final score for each residue to a maximum of 2. The weighted match scores are then summed across interfacial residues to give the final Nanohedra score for the identified docking configuration.

### Program considerations

Nanohedra is a command line tool. It can be operated in one of three modes: Query, Dock or Post-processing. The Docking mode executes the main procedures described in the present work. The user specifies the desired symmetry material outcome or SCM type, i.e. the specification of the two component symmetries and their resulting assembly type. Directory paths are input to specify the file locations for the oligomeric protein structures to be tested. The program output comprises pdb files with candidate docked poses in various forms (asymmetric unit within the final symmetry, docked oligomers, and an expanded symmetry). Other information includes the final Nanohedra score and the spatial transformation matrices mapping the canonically oriented coordinates onto the candidate pose. To guide subsequent design of the resulting interface, sequence information is output in the form of amino acid frequencies based on amino acid composition information tabulated from the fragment database (Fig. S3).

The computer time for execution depends critically on the size of the proteins (because larger proteins carry more surface fragments), the number of rotational degrees of freedom for sampling, and the rotational sampling interval. Times on a single CPU core (2.5 GHz) can range from 2 to 24 hours, with typical applications exploiting multi-core clusters. Computer memory requirements also depend on the sizes of the proteins and the size of the symmetry group generated by the final assembly. Requirements range from roughly 8 to 25GB. The user can override various default settings, e.g. angular sampling in rotational searching (-rot_step1/-rot_step_2) or the minimum number of fragment-fragment pairs needed for a well-docked pose (-min_matched).

Query mode is an informational mode that helps the user understand different options and certain symmetry aspects of the material to be designed: e.g., what kinds of resulting SCM materials can be constructed from a given combination of components, and conversely what component oligomer types would be needed to construct different SCMs according to various target criteria, such as the dimensionality of the resulting material (cage vs layer vs three-dimensional crystal), the underlying rotational symmetry, or specific geometric features (like network properties) of the material to be designed. Different materials properties will be advantageous in different experimental contexts, and this mode captures the full space of design types recently articulated [40]. A final Post-processing mode provides tools for ranking the output candidate poses, with options to sort by different criteria, e.g. according to the final Nanohedra matching score or according to the numbers of fragment pairs identified in the match.

The program is implemented in Python with the exception of one routine that is written in Fortran (orient_oligomer). Python dependencies include biopython [42], numpy [43], and scikit-learn (the BallTree method is used to test for clashes) [44]. Nanohedra also uses the freeSASA program to calculate solvent accessible surface areas [45]. The program code has been made available on GitHub.

### Prospective SCMs

To demonstrate the universality of our fragment-based docking approach, we constructed prospective SCMs of six divergent types, with representatives from point (1), layer (2), and space group symmetries (3), based on component oligomer symmetry types ranging from C3 to T. Nanohedra was run with default docking parameters, and for each SCM type a search of the rigid body degrees of freedom inherent in each system produced numerous viable candidates with varied orientations and positions and different interfacial secondary structure compositions. Post processing revealed numerous poses with high Nanohedra scores. One representative structure produced for each of the six SCM types tested is displayed in Figure 3. The results demonstrate the viability of the described method at producing assemblies conforming to a selected symmetric material.

**Fig. 3.**
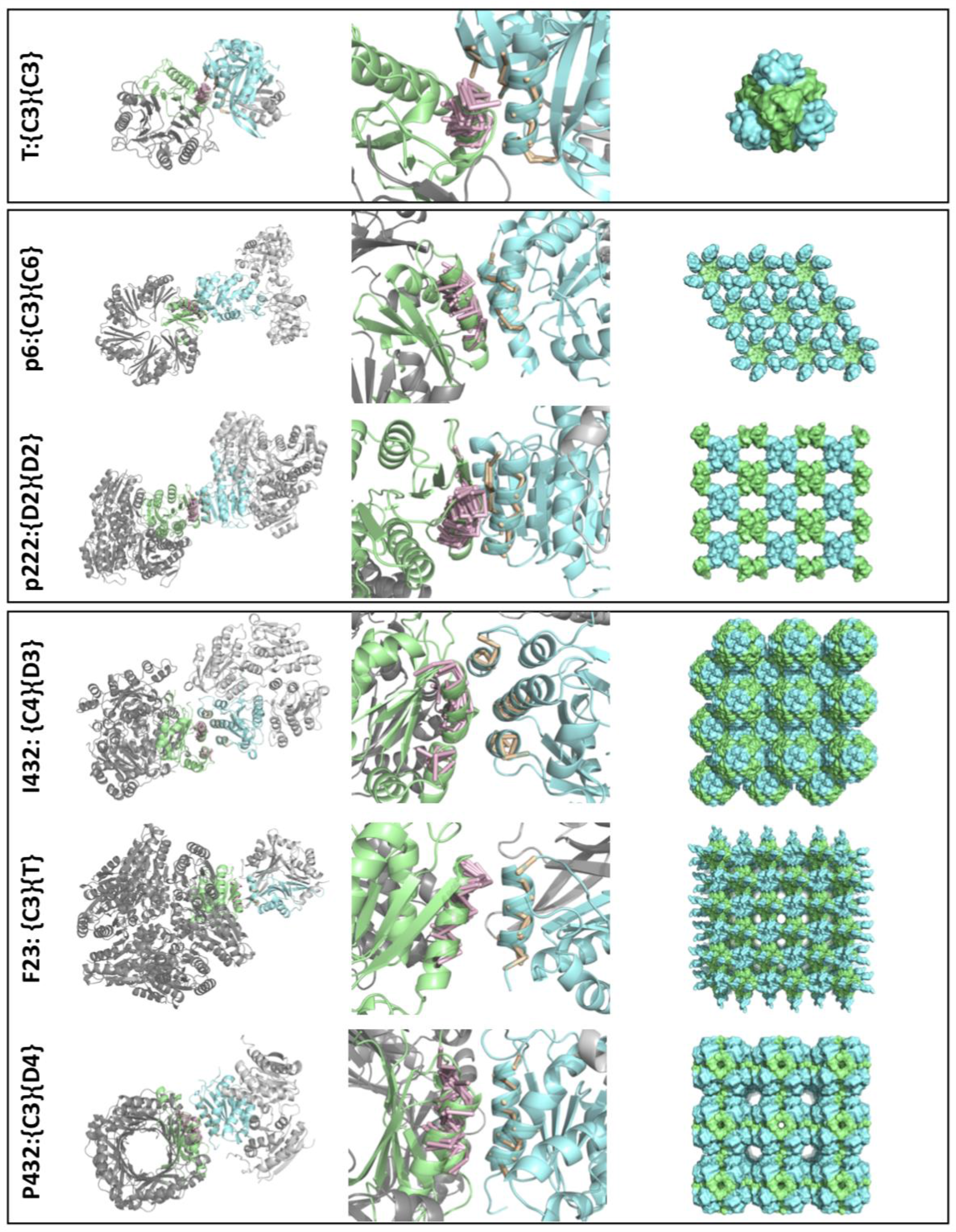
Prospective SCMs. Six example SCMs generated by Nanohedra are shown: a finite tetrahedral cage (top), two 2-D layers (middle), and three 3-D crystals (bottom). Column 1 illustrates the docked oligomeric building blocks that are required to construct the final material. Chains directly implicated in the docked interface are colored, while symmetrically related chains are in grey. Closeups of the interfaces are shown in column 2. Oligomer 1 (blue) surface fragments (tan) and associated ghost fragments (pink) are shown. Ghost fragments are matched with secondary structure elements on the surface of oligomer 2 (green). The resulting symmetrically expanded materials are displayed in column 3. PDB accession codes used to generate the prospective materials shown are: 1OSC and 1NQ3 for T: {C3}{C3}, 4O5O and 1UAY for p222: {D2}{D2}, 1OSC and 1GTZ for F23: {C3}{T}, 2B34 and 3BBC for I432: {C4}{D3}, 1VHC and 2A10 for p6: {C3}{C6}, 4XCW and 1DHN for P432: {C3}{D4}.

In each example, the resulting interface exhibits native-like properties with respect to interfacial backbone-backbone associations. High structural complementarity between oligomers is apparent from the overlap between the ghost fragments of the first oligomer and the matched surface fragments of the second oligomer. The interfaces vary in the extent of regular secondary structure involvement, with each oligomer contributing at least one continuous secondary structure element to the interface, and ranging from 8 (F23:{C3}{T}) to 20 (p222:{D2}{D2}) unique fragment matches (Fig. 3). Many of the docked configurations comprise extensive helical interactions, with interfaces containing anywhere from two to five helices (see F23:{C3}{T} and I432:{C4}{D3}, respectively). The contribution from β-strands is also apparent as both T:{C3}{C3} and p222:{D2}{D2} designs have mixed α/β interface motifs, despite prioritizing helix-helix pairs in first-pass searching. Additionally, matched interface fragments are sometimes enhanced by fortuitous contacts involving coiled segments surrounding regular secondary structures. These characteristics are reminiscent of patterns observed in nature [31].

### Post facto analysis of designed protein cages

Prior work in designing protein assemblies has shown the challenges of generating computational designs that produce the desired experimental outcomes; success rates remain relatively low, as failures can manifest at many crucial junctions. The favorable features of Nanohedra—constructing designs based on native-like interfacial packing—will ultimately require experimental tests that are ongoing and not presented here. Nonetheless, the results of several recent design trials provide an opportunity to evaluate the prospective advantages of Nanohedra ahead of new experimental trials.

For a retrospective analysis, we asked whether Nanohedra could distinguish experimentally validated designed protein assemblies among a larger body of prospective computational designs that were unsuccessful. We focused on designed protein cages, for which there are more than a dozen successful cases validated in atomic detail, along with more than a hundred computational designs that led to experimental failure. We ran Nanohedra on these designs to see if there was a difference in the generation of candidate poses that matched prior design targets between the two sets; this would argue that Nanohedra has the capacity to generate computational designs that have improved experimental success rates. For each prior design (in both categories of experimental successes and failures), we took the two component oligomers in standard orientations (i.e. not corresponding to the previously designed configurations) and ran Nanohedra to generate prospective designs for symmetric cages of the desired symmetry. While Nanohedra was able to recapitulate the target in nearly all cases, there were differences in the extent to which the design target was ranked favorably compared to other potential designs comprising the two oligomers. We clustered all poses in the top 2000 output, then examined the ranked output to see where in the list of candidate poses (if at all) we could find configurations closely matching the target that was experimentally tested in earlier work. For the group of experimental successes (n=14), we were able to recapitulate 70% of the design targets within the top 12 scoring pose clusters for each combination of building blocks and 100% of the targets within the top 88 ranked poses (Fig. 4). In contrast, for the unsuccessful design set (n=138), we were unable to identify closely matching poses for 25% of designs, and had to search until rank 112 in order to recapitulate 70% of the designed poses. These calculations clearly show that among earlier computational designs, those that went on to experimental success are much more readily recapitulated using Nanohedra compared to designs that failed. This indicates that using motifs present in native interfaces leads to improved search heuristics for biologically confirmed symmetric material designs. We further note that having to go to rank 12 to recapitulate most of the earlier experimental successes does not preclude that poses ranked higher by Nanohedra could quite plausibly lead to successful experimental constructions, different in orientation from those validated earlier.

**Fig. 4.**
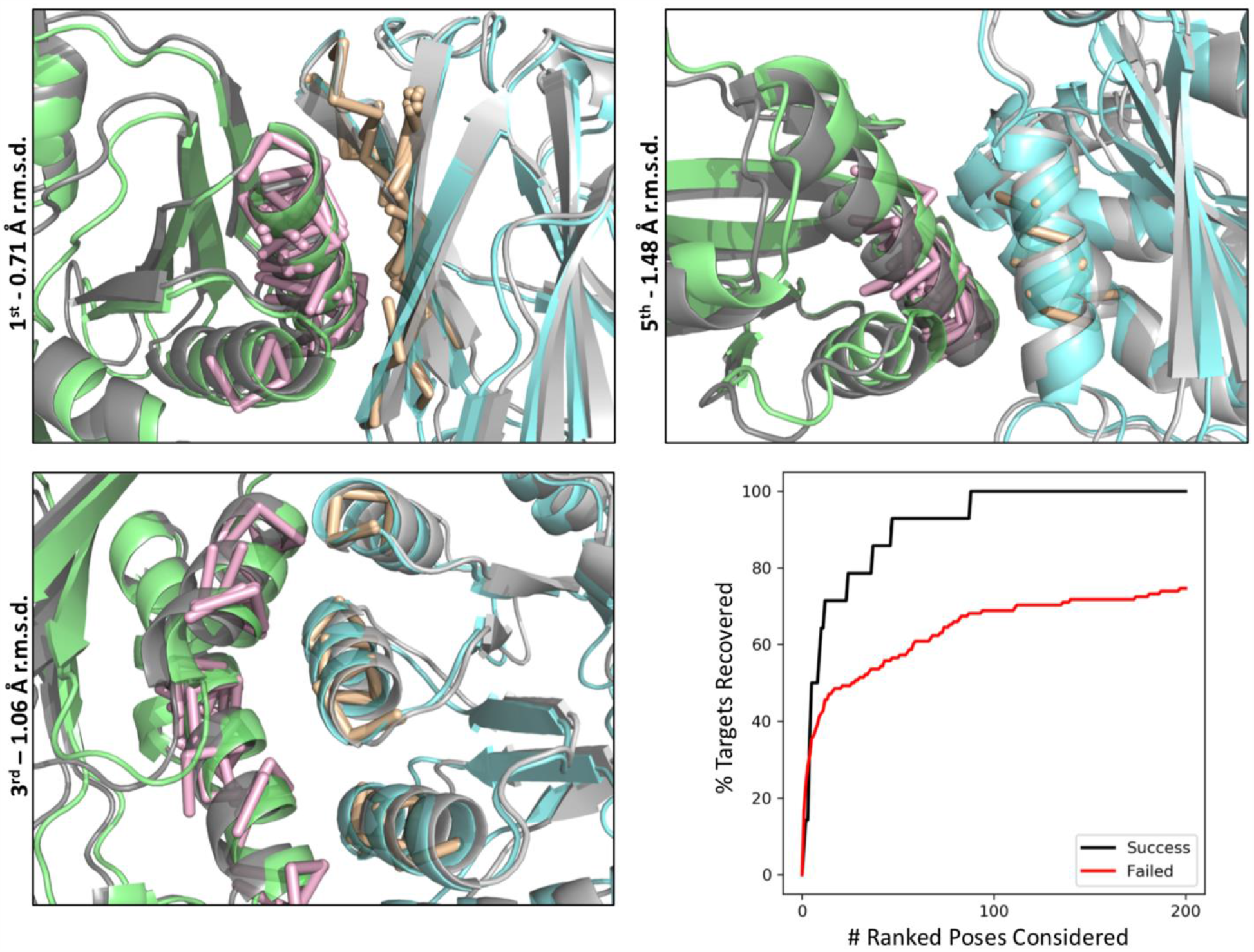
Post facto analysis of designed protein cages. Three representative examples of successfully recapitulated poses (top and bottom left). The crystal structures of the target designs are shown in grey and Nanohedra predictions are shown in color. The iRMSD indicates the agreement, for atoms near the interface, between the docked pose and the crystal structure. The numerical value indicates the rank of the docked pose. The crystal structure of T32-28 (4NWN) displays a 0.71 Å iRMSD with the first ranked pose (top left). The crystal structure of T33-15 (4NWO) exhibits a 1.48 Å iRMSD with the fifth ranked pose (top right). The crystal structure of I53-40 (5IM5) shows a 1.06 Å iRMSD with the 3rd ranked pose (bottom left). The ROC curve for all successful and failed designs shows the percentage of targets recovered according to the number of clustered Nanohedra poses considered (bottom right). iRMSD of 3.0 Å or less was considered as a recovered pose. iRMSD - interface root mean squared deviation.

## DISCUSSION

Until now, designing symmetric protein assemblies has remained challenging to new entrants, as the process requires somewhat expert knowledge about symmetric construction and intertwined issues of how to sample allowable degrees of freedom in the context of docking software. Previously successful studies in executing two-component symmetric docking have used the Rosetta TCdock protocol, which requires the user to specify the symmetry rules through the use of symmetry definition files [4]. This is possible for symmetries already enumerated, but the majority of recently described SCM’s [40] present a remaining challenge for specifying allowable degrees of freedom in the context of existing design software. By enumerating the allowable degrees of freedom for each SCM in a comprehensive, facile framework, Nanohedra will empower a broader group of users to explore the large space of possible symmetric designed materials. This should accelerate the development of novel designed materials by protein and biomaterial engineers.

Nanohedra harnesses the power of a recently established theoretical framework [40] to enable the construction of a universe of possible protein-based nanomaterials. In addition, Nanohedra’s docking algorithm implements a novel fragment-based approach for assessing whether docked solutions resemble biological interfaces. Importantly, the Nanohedra score does not depend on commonly used heuristic simplifications (such as number of Cβ-Cβ contacts) for rapid assessment of binding likelihood, as has been used in various docking studies [4,5,22]. Instead, its statistical representation of clustered secondary structure elements exploits empirical knowledge of typical packing motifs found in native protein interfaces. The implications of this choice, as demonstrated in our retrospective analysis of successful versus failed designs, are notable. In agreement with previous findings, geometric packing alone captures many essential elements of protein interaction [46]. Furthermore, although Nanohedra is geared for design of symmetric materials, our results point to potential opportunities for advances in macromolecular docking and interface design in other contexts.

This first version of Nanohedra will admit future improvements along various lines, including GPU enhancements to increase speed. Critical experimental studies will be needed to evaluate the most important assertions concerning the expected advantages of generating assemblies with native-like interfaces. We also emphasize that successful design requires judicious amino acid interface design as a final step. This subsequent design step is separate from the construction of native-like (backbone-level) poses provided by Nanohedra, though as noted above, Nanohedra provides valuable information about specific amino acid preferences favored in the fragment interfaces. This position specific frequency information can be exploited by sequence design programs in the final step of design.

## METHODS

### Fragment database generation

To generate the fragment library, all non-redundant, biologically relevant interfaces from high resolution structures were gathered from the PDB. For homomers, structure codes for biological assemblies referenced in the QSBio database [47] were used to extract all assemblies that were verified with a confidence ranking of ‘high’ or ‘very high’. For heteromers, biological assemblies were identified using PISA [48]. The homo- and heteromer sets were next filtered to include only representative structures clustered at 90% sequence identity with a reported resolution <= 2.0 Å, experimental expression in *E. coli*, no nucleic acids and no membrane proteins. For each identified structure, all unique interfaces between two separate chains were extracted excluding chains with less than 10 residues or fewer than 5 Cβ atoms within 8 Å of a second chain. From the resulting chain pairs, inter-chain Cβ distances were computed and residues that were 8 Å or less apart were selected as residue pairs across the interface. For each residue in the interface residue pair, the preceding and following two residues (i.e. i-2 through i+2) were included in the observation and the resulting 5-residue segments were stored, first as an individual 5-residue segment (individual fragments), and second as a pair of 5-residue segments across the interface (paired fragments). For residues with multiple conformations, the A conformation was chosen. Selenomethionine residues were not considered.

From the pool of individual fragments, a subset was chosen to perform all-against-all RMSD measurements followed by nearest neighbor clustering. The top five neighbor clusters were selected as the clustering population significantly decreased after this point. From each of the top five clusters, the fragment with the most neighbors was selected as a cluster representative, centered on the origin, and stored. Each of these five clustered fragments represents one unique type of individual fragment, and the instance with the most neighbors was chosen as the fragment representative. For the saved paired fragments, both fragments in the pair were queried for membership in one of the five individual fragment types according to a Cα RMSD threshold of 0.75 Å. If one of the fragments in the pair did not belong to an individual fragment type, the pair was discarded from further classification. Next, each fragment in the fragment pair was subjected to a structural superimposition on its corresponding matched individual fragment representative. This centered one fragment in the pair at the origin aligned to its structural representative, while maintaining the relative position of the partner fragment to this aligned fragment. Once in this orientation, a set of three guide coordinates was stored, one coordinate at the partner fragment’s central Cα atom, the second displaced by a unit vector along the C-alpha to subsequent carbonyl carbon vector, and the third displaced by a unit vector perpendicular to the previous vector and lying in the plane formed by the C-alpha atom, the subsequent carbonyl carbon and the preceding amide nitrogen. This guide coordinate set, stored for each fragment observation, describes the transformation of the partner fragment’s central Cα atom, and its relative orientation with respect to the aligned individual fragment representative. In this way, each partner fragment provides a unique spatially encoded and secondary structure dependent observation of the interaction potential surrounding each individual fragment type.

Finally, for each individual fragment representative, and for each set of secondary structure dependent guide coordinates of that fragment representative, a subset of those guide coordinates was subjected to all against all RMSD calculations followed by nearest neighbor clustering. The resulting guide coordinate clusters were binned with a maximum of 1 Å deviation, requiring at least four members in the cluster to be considered. From this set of guide coordinate clusters, all possible guide coordinates were subjected to membership in the resulting clusters by testing for the minimal RMSD to an established cluster. If a cluster with RMSD less than 1 Å could not be located, the guide coordinates were disregarded as outliers. This procedure was applied for each partner secondary structure associated with each fragment representative.

For each i,j,k fragment pair cluster, the cluster representative fragment coordinates and guide coordinates were stored. Additionally, the cluster size, mean guide coordinate RMSD and observed amino acid pair frequencies for central fragment residues were stored. The top 75% most populated i,j,k clusters were then chosen for our final fragment database.

### Docking prospective SCMs

From the set of 124 possible SCM types, we chose six as diverse representatives for presentation in this study; note that all 124 were tested for mathematical and computational correctness in our earlier study [40]. For each of these SCM types, homo-oligomers matching the design criteria were curated from the PDB by searching for the desired point group symmetry, X-ray resolution better than 2.5 Å, a helical content greater than 30% and *Escherichia coli* as the organism used for protein expression. Structures containing membrane proteins or nucleic acids were removed. Biological assemblies were identified using QSBio [47] and representatives clustered at 70% sequence identity were then downloaded from the Protein Data Bank. A few candidate oligomeric building blocks were then selected for pair-wise docking with Nanohedra using the default parameters.

### Design recapitulation

The dataset for the design recapitulation experiments was generated by selecting all successfully designed two-component tetrahedral, octahedral and icosahedral designs from previously published work [4–6,12,14,49] ; these cases met the criteria of agreement between the model and an experimentally determined atomic model. Failed designs (e.g. described as insoluble or unknown oligomerization state) were also identified from earlier studies [4,5].

For each successful design, the two component oligomers used for docking were extracted from the deposited PDB structure of the protein cage. For the failed designs, the PDB structures of the native oligomeric building blocks were used. Default Nanohedra docking parameters were used with the exception of a 2° rotational sampling step instead of 3° for each component oligomer. Docking proceeded until all rotational degrees of freedom had been sampled. For 4NWN we had to modify the initial default Helix-Helix fragment search to Strand-Helix. Since the dimeric component is mainly composed of β-strands on its surface, suitable docked configurations could not be identified with the default initial Helix-Helix search. Only the default Helix-Helix fragment search was used for failed designs, and designs were not considered in rare cases where no helix-helix interaction was present.

The Cα interface RMSD (iRMSD) was computed between the target design and each Nanohedra output pose. For successfully designed structures, the coordinates deposited in the PDB were used as a reference. Models of the failed designs noted in earlier studies [4,5] were obtained from Neil King and Jacob Bale. For each design target, the 2000 docked poses with the lowest iRMSD to the design target were selected and nearest neighbor clustering was performed using all to all iRMSD calculations. Interfaces within 1Å iRMSD threshold were clustered, then each cluster was ranked according to the Nanohedra score of the cluster representative.

### Amino acid frequency plots

The Nanohedra program outputs amino acid frequencies for the central residue in the docked pose for each surface-ghost fragment match that has been identified. To calculate this frequency distribution, frequencies are retrieved from the fragment database for the corresponding i,j,k cluster for each surface-ghost fragment pair. When multiple surface-ghost fragments are identified for the same residue, the frequency distribution is a sum of the individual amino acid frequencies, proportionally weighted by the corresponding surface-ghost fragment match score. In this instance, the final distribution reflects the separate constraints of all identified fragments. Weighting the frequency distribution in this way provides a quantitative output for how well the amino acid identities from the fragment library fit within the specified docked conformation. To visualize these distributions, at each residue the resulting frequencies were transformed into multiple sequence alignments and sequence logos were generated using the WebLogo server [50].

## CODE AVAILABILITY

The Nanohedra source code is freely available at https://github.com/nanohedra/nanohedra

## ACKNOWLEDGEMENTS

This work was supported by NSF Grant: CHE-1629214. KM was supported by NIH training grant T32GM008496. We thank Duilio Cascio and Alex Lisker for computing support and Michael Sawaya for helpful discussions. We thank Neil King and Jacob Bale for providing us with the models of their two-component cage designs.

## AUTHOR CONTRIBUTIONS

The research was conceived by TOY and JL. The code was written by TOY and JL. The fragment database was constructed by JL. The example SCMs were constructed by JL and KM. The post facto analysis of designed protein cages was performed by KM and JL. The manuscript was prepared by TOY, KM and JL.

## NOTES

The authors declare no conflicting interests.

