## Supplementary Materials for "A fragment-based protein interface design algorithm for symmetric assemblies"

**This PDF file includes**

Supplementary Text

Supplementary Figures S1 – S3

**SUPPLEMENTARY FIGURES**

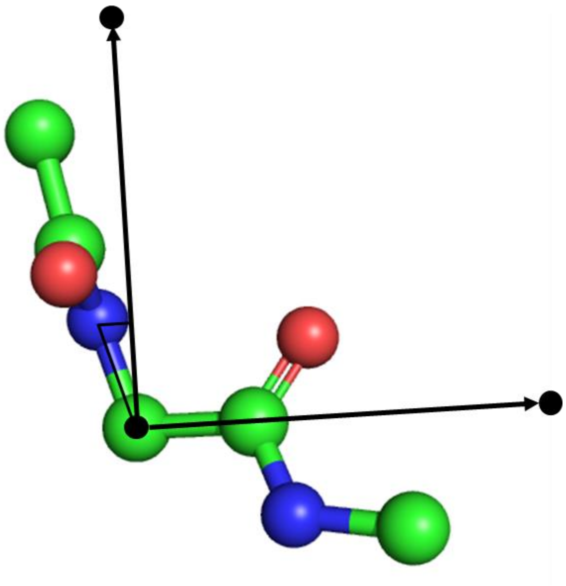

**Fig. S1.** Guide coordinates. An orthonormal three-atom system is constructed on the central C-alpha position of a 5-residue fragment to provide a reduced representation of its position and orientation. The unit vector length for typical calculations in Nanohedra is set to 10 Å.

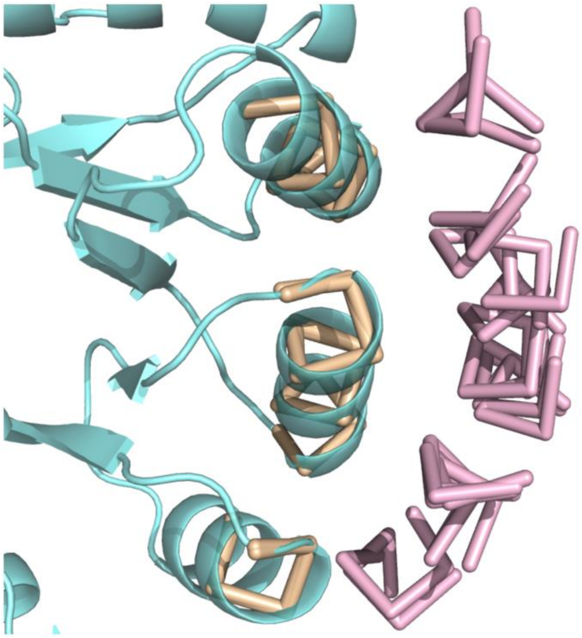

**Fig. S2.** Example illustration of ghost fragments. Oligomer 1 (cyan) is shown with its surface fragments (tan) and associated ghost fragments (pink), which represent candidate fragment associations.

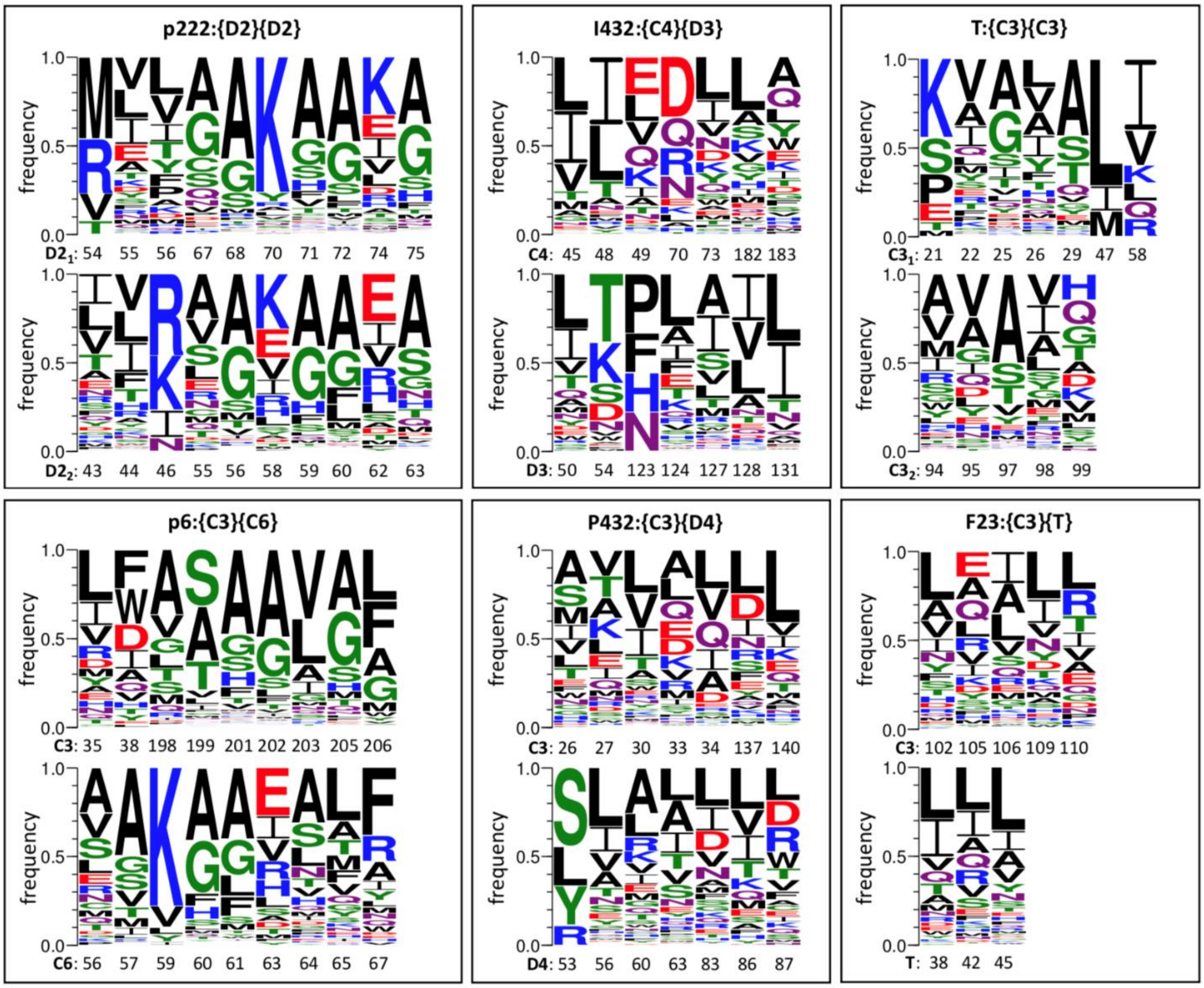


**Fig. S3.** Deduced amino acid preferences for prospective SCMs in Figure 3. For each case, the top and bottom diagrams indicate the preferences for the two oligomeric components, whose symmetry types are noted to the left of the participating residue numbers in the interface.

### SUPPLEMENTARY TEXT

<u>PDB IDs and design names used for protein cage design recapitulation experiments</u>

PDB IDs of the experimentally validated designs:
4NWN, 4NWO, 4NWP, 4NWR, 4ZK7, 5CY5, 5IM4, 5IM5, 5IM6, 6P6F, 6VFH, 6VFI, 6VFJ, 6VL6

Names of the 'failed' designs:
I32-01, I32-03, I32-05, I32-07, I32-08, I32-12, I32-13, I32-14, I32-15, I32-16, I32-17, I32-20, I32-22, I32-23, I32-24, I32-25, I32-27, I32-31, I32-33, I32-34, I32-35, I32-36, I32-37, I32-38, I32-39, I32-40, I32-41, I32-45, I32-46, I32-49, I32-52, I32-53, I32-54, I32-55, I32-56, I32-60, I32-62, I32-

64, I32-65, I32-68, I32-70, I52-01, I52-07, I52-09, I52-10, I52-11, I52-12, I52-14, I52-17, I52-18, I52-20, I52-22, I52-23, I52-24, I52-26, I52-27, I52-28, I52-29, I52-31, I52-34, I52-35, I52-36, I52-38, I52-39, I52-40, I52-41, I52-42, I52-43, I52-44, I52-46, I53-06, I53-09, I53-12, I53-15, I53-16, I53-21, I53-23, I53-25, I53-27, I53-28, I53-29, I53-33, I53-35, I53-37, I53-43, I53-48, I53-49, I53-58, I53-61, I53-62, I53-63, I53-64, I53-67, I53-68, I53-70, I53-72, I53-74, I53-75, I53-79, I53-80, I53-82, I53-83, T32-02, T32-03, T32-06, T32-07, T32-08, T32-09, T32-11, T32-17, T32-18, T32-20, T32-24, T32-25, T32-26, T32-27, T33-01, T33-02, T33-03, T33-04, T33-05, T33-06, T33-07, T33-08, T33-11, T33-12, T33-13, T33-14, T33-16, T33-17, T33-18, T33-22, T33-23, T33-24, T33-25, T33-26, T33-27, T33-29

RPX designs were those utilizing the motif library from [1] which included I32 and I52 designs. All others are Non-RPX designs.
